# Genome urbanization: Clusters of topologically co-regulated genes delineate functional compartments in the genome of *S. cerevisiae*

**DOI:** 10.1101/064667

**Authors:** Maria Tsochatzidou, Maria Malliarou, Nikolas Papanikolaou, Joaquim Roca, Christoforos Nikolaou

## Abstract

The eukaryotic genome evolves under the dual constraint of maintaining co-ordinated gene transcription and performing effective DNA replication and cell division, the coupling of which brings about inevitable DNA topological tension. DNA supercoiling is resolved and, in some cases, even harnessed by the genome through the function of DNA topoisomerases, as has been shown in the concurrent transcriptional activation and suppression of genes upon transient deactivation of topoisomerase II (topoII). By analyzing a genome wide run-on experiment upon thermal inactivation of topoII in *S.cerevisiae*. we were able to define 116 gene clusters of consistent response (either positive or negative) to topological stress. A comprehensive analysis of these topologically co-regulated gene clusters revealed pronounced preferences regarding their functional, regulatory and structural attributes. Genes that negatively respond to topological stress, are positioned in gene-dense pericentromeric regions, are more conserved and associated to essential functions, while up-regulated gene clusters are preferentially located in the gene-sparse nuclear periphery, associated with secondary functions and under complex regulatory control. We propose that evolves with a core of essential genes occupying a compact genomic “old town”, whereas more recently acquired, condition-specific genes tend to be located in a more spacious “suburban” genomic periphery.

## INTRODUCTION

The distribution of genes in the genome of eukaryotes is highly non-random. Early genome-wide transcriptome analyses showed the expression of genes to correlate with their linear order along the genome (1). Although it was later shown that this was due to the clustering of constitutive genes (2), such spatial associations have since been used to provide the theoretical framework for links between gene expression and chromatin structure (3) and the inference of protein-protein interaction patterns (4). Non-random gene distribution is also evident in the functional enrichments of gene neighborhoods, with functionally related genes being found in linear proximity more often than expected by chance (5,6).

The selective pressures underlying gene localization are thus of unequal intensity and diverse nature and a number of seemingly irrelevant characteristics may shape the overall genome architecture through evolution (7). Among those DNA supercoiling plays a prominent role. The structure of the eukaryotic nucleus is affected by a number of processes such as DNA replication, RNA transcription and the constant ebb and flow of gene activation and repression. These processes are imposing topological constraints in the form of supercoiling, both types of which (positive and negative) may be found in localized areas of the eukaryotic genome (8). It was recently shown that such structurally-defined areas may form part of extended “supercoiling domains”, where chromatin conformation correlates with the density of topoisomerases I and II (9). The connection between topological attributes and gene expression appears to be so strong, that in *Drosophila melanogaster* regions of negative supercoiling, created through the inhibition of topoisomerase I, show increased nucleosome turnover and recruitment of RNA-PolII molecules positively correlating with transcription levels (10). Accumulated positive supercoiling, on the other hand, precludes the formation of transcription initiation complexes (11,12), a fact indicative of the association between topological constraints and gene expression.

In the budding yeast (*Saccharomyces cerevisiae*), the organization of genes in linear space has also been attributed to common regulatory mechanisms (13). Yeast’s distinguishing genomic feature among eukaryotes, is the overall gene density, with genes covering ~70% of the total genome (14). Despite its reduced size of only 12Mbp, the transcription dynamics of the yeast genome is highly complex, with genes being expressed in tandem and in operon-like transcripts, with varying sizes of gene upstream and downstream regions (15). Transcription directionality in such a highly streamlined genome also plays a crucial role in the regulatory process, with a number of bidirectional promoters (16) exerting control over coupled gene pairs. The interplay between DNA structure and gene regulation is manifest in a number of cases where gene expression is modulated through three-dimensional loops formed at gene boundaries (17). Thus, even in a small eukaryotic genome, there is a strong association between gene organization (in both linear and three-dimensional space) and gene expression.

The response to topological stress has been shown to be shaped by specific structural properties of yeast promoters (18). In this work, we sought to investigate how the response to the accumulation of topological stress may extend beyond single gene promoters to affect broader genomic regions. Starting from a Genomic transcription Run-On (GRO) experiment, we explored the formation of clusters of genes that are differentially affected by topoII deactivation and then went on to assess a number of related functional and structural preferences. We were able to detect intricate associations between DNA topology and the distribution of genes in linear order and to show how the two may be linked to other organizational characteristics such as gene spacing, transcriptional directionality and the three-dimensional organization of the yeast genome. Our results are suggestive of a subtle dynamics of evolution of genome architecture, which we describe as “Genome Urbanization” and according to which the relative position of genes in the nucleus reflects a broader functional, structural and regulatory compartmentalization.

## MATERIALS AND METHODS

### GRO data

Data were obtained from a genome-wide Genomic Run-On (GRO) experiment conducted in triplicates on a yeast strain lacking topoisomerase I and carrying a thermosensitive topoisomerase II (JCW28 - top1Δ, top2ts). GRO was conducted as described in (19) and data were analyzed as previously described in (18).

### Gene Clustering

Starting from an initial dataset of differential GRO values for 5414 yeast protein coding genes (Supplementary File 1), gene clusters were defined as the uninterrupted regions spanning the genomic space from the first to the last segment in an all-positive (up-regulated) or all-negative (down-regulated) gene series (Figures 1A,B). Clusters of >=7 genes were selected on the basis of a bootstrapping analysis as suggested in (7). This was performed by conducting 10000 random permutations of gene order while keeping the same GRO values. We used functions from the BedTools Suite (20) to control for unaltered gene sizes and chromosomal distributions. Gene number distributions of the derived clusters were calculated alongside the mean values and standard deviation of number of clusters for the 10000 random gene sets. We then compared the observed values with the expected under randomness asking that the observed value be at least greater than the mean of the 10000 permutations by two standard deviations. Clusters with >=7 genes occurred in less than 0.1% of the simulations (bootstrap value p=0.0008) and were divided into up-regulated and down-regulated, depending on the mean GRO value of all genes in each cluster (Supplementary File 2).

**Figure 1.**
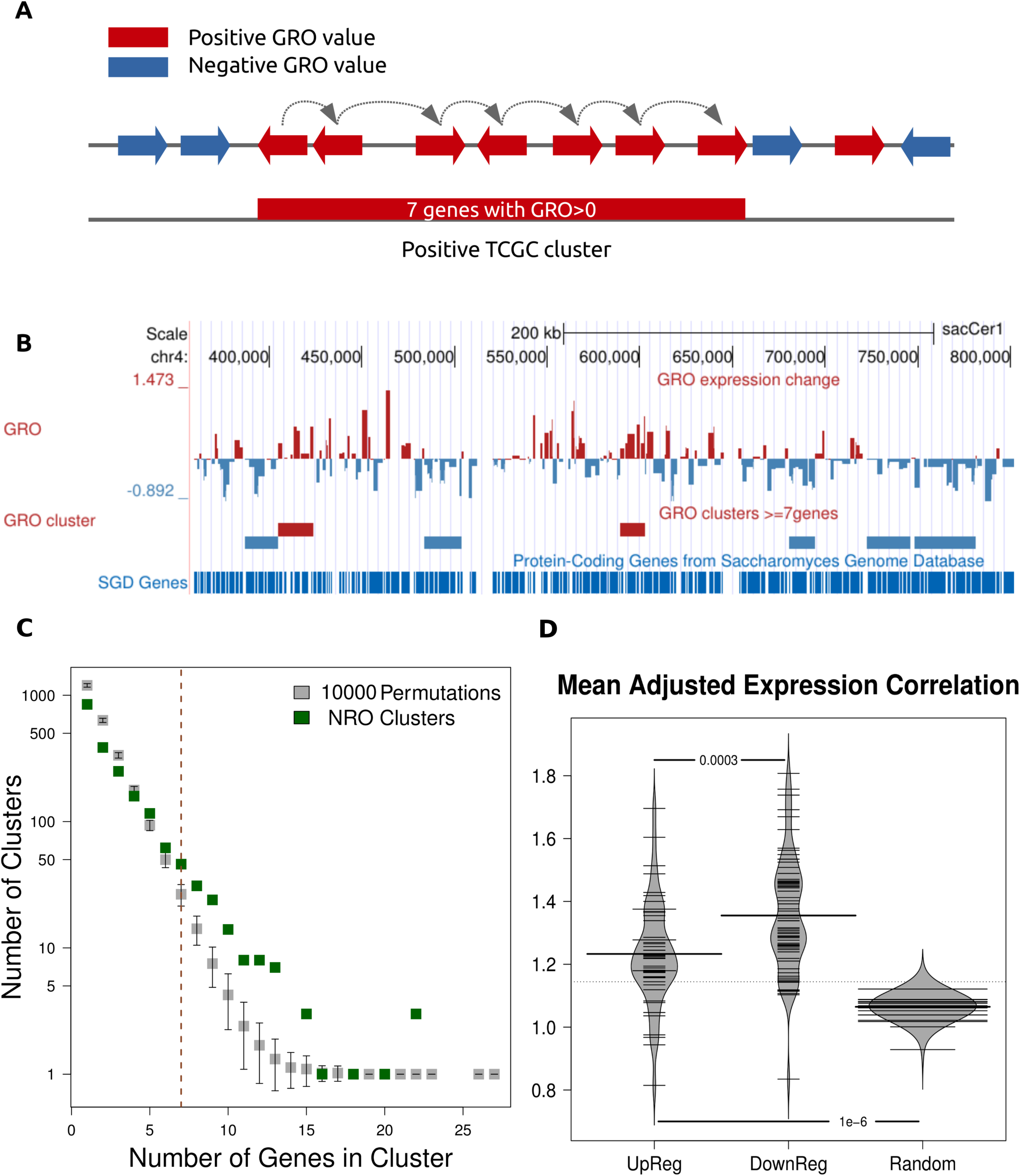
DNA topological stress-responsive genes are clustered non-randomly. **A.** Schematic representation of cluster calls. Contiguous genes with similar (positive or negative) GRO values were joined in gene clusters, which were defined as the genomic region spanning the chromosomal space from the fartherst upstream to the farthest downstream gene boundary. **B.** Location of genes and clusters with GRO values in part of chromosome 4. **C.** Distribution of number of genes in clusters. Real clusters (green) show a skewed distribution towards larger sizes as compared to the mean of 10000 random permutation of GRO values. Differences are significant for gene numbers >=6. **D.** Mean Adjusted correlation scores (ACS) for up– and down-regulated gene clusters are significantly higher than the genomic average (randomly selected clusters of equal size and number), indicating increased co-regulation within the confines of the defined clusters.

### Gene Cluster co-expression Index

Adjusted Correlation Scores (ACS) were obtained for the complete set of yeast genes from the SPELL Database (21). ACS values represent weighted correlation values for a large number of genome-wide expression profiles. As a measure of co-expression in a gene cluster, we calculated the mean ACS of all genes within the confines of the cluster.

### Positional enrichments of gene clusters in one dimension

Genomic coordinates for yeast centromeres were obtained from SGD (http://www.yeastgenome.org/locus/centromere). Cluster-centromere distances were calculated as the sequence length between the most proximal cluster boundary to the central point of the centromeric coordinates. Distances were then scaled with the size of the chromosomal arm extending from the central point of the centromere to the chromosome’s boundary, so as to be represented in a range of 0 (i.e. overlapping the centromere) to 1 (i.e. lying at the edge of the corresponding chromosomal arm). Distances from Autonomously Replicating Sequences (ARS) were calculated in the same way based on a compiled list of 829 yeast ARS published in OriDB (http://cerevisiae.oridb.org/) (22).

### Three-dimensional positional enrichments of gene clusters

We obtained the raw frequency measurements of a yeast 3C experiment (23). In order to define TAD-like domains, we used the insulation profile approach described in (24), where an aggregate score of contact frequencies is calculated along the diagonal of an interaction map. By setting an upper limit of insulation score equal to the bottom 5%-percentile we were able to define 86 insulation domains at 10kb resolution. We then compared these domains for overlaps with the defined gene clusters.

At a second level we used the classification of yeast chromosomal regions in network communities described in (25). We calculated the enrichment of our gene clusters, separately for up– and down-regulated ones in the 13 distinct level-1 communities (Supplementary Table 7 from (25)). Enrichment was calculated on the basis of an observed over expected ratio of overlaps between the two sets of genomic coordinates and was statistically assessed on the basis of 1000 random permutations of cluster coordinates. Overlaps with a bootstrap p-value less or equal to 0.01 were deemed significant (Supplementary File 3).

### Functional and Regulatory Enrichment

We employed a modified gene set enrichment functional analysis at gene cluster level to analyze concerted over-representations of Gene Ontology terms (www.geneontology.org). Enrichment was calculated based on a hypergeometric test for each gene cluster and controlled for multiple comparisons at a 5% FDR (26). GO terms with significant enrichment (adjusted p-value <=0.05) in at least one of the two types of gene clusters (up– or down-regulated) were recorded.

Conserved Transcription Factor Binding Sites (TFBS) were obtained from the UCSC Genome Browser’s Transcriptional Regulatory Code track. These corresponded to a compendium of 102 transcriptional regulators based on a combination of experimental results, cross-species conservation data for four species of yeast and motifs from the literature initially compiled by (27) and updated by (28). Enrichment in TF binding was calculated as in the case of chromosomal communities described above. Enrichments were assessed as ratios of observed over expected overlaps and p-values were obtained as bootstrap values from 1000 random permutations of cluster coordinates.

### Gene and intergenic space size and direction of transcription

We used genomic coordinates downloaded from UCSC (SGD/saCcer2). Intergenic distances were calculated as the full length of regions spanning the genomic space between two consecutive genes, using transcription initiation and termination as boundaries, regardless of gene transcription direction. We assigned to each gene a mean intergenic space length to be the arithmetic mean of the lengths of gene upstream and downstream intergenic regions. For genes at chromosomal boundaries, one of the two intergenic regions were set to be equal to the distance from the gene boundary to the corresponding chromosomal start/end.

For the analysis of gene directionality, each chromosome was scanned in overlapping 11-gene windows and for each step we recorded: the full list of 11 GRO values, mean GRO value of the central 7 genes and gene lengths and mean intergenic space lengths for all genes. The top/bottom 200 non-overlapping clusters in terms of mean GRO value were analyzed at the level of gene and intergenic spacer lengths (Figure 3A). We used the same list to obtain patterns of gene directionality as arrays of seven genes (Figure 3B). GRO values of the central gene were analyzed for three characteristic patterns corresponding to a) co-directional genes (central 5 genes transcribed in the same direction) b) the central gene being a member of a divergent or c) a convergent gene pair.

### Sequence conservation and TFBS density

Sequence conservation was calculated as aggregate phastCons scores (29) obtained from UCSC and based on a multiple alignment of 7 *Saccharomyces* species. Mean conservation was taken as the mean phastCons score for a given region. For each cluster we removed intergenic space and calculated the mean aggregate phastCons score for all genes in the cluster. TFBS density was calculated as the percentage of the length of each TCGC overlapping with conserved TFBS as compiled in (27).

### Gene Cluster Directionality Conservation Index

We obtained orthologous gene coordinates for *S. paradoxus* and *S. mikatae* from the Yeast Gene Order Browser (YGOB) (30). For each genomic region of *S. cerevisiae* we calculated the ratio of genes retaining their position and direction of transcription in the other two species. A value of 1/N, N being the number of genes in the region, was added to the score if both the gene’s position and direction was maintained in the other two species. This led to measure of directionality conservation on a scale of 0 (no retention of direction) to 1 (absolute retention of direction). The contour map of Figure 3C was formed by splitting the two-dimensional space in a 10x10 grid and assigning each bin with the proportion of clusters falling in the corresponding sequence/direction conservation value range (bins of 0.1 for each). The final value assigned to each of the 10x10 bin was the log2(ratio) of up/down-regulated cluster frequency. Values >0 corresponded to an enrichment of up– and values < 0 to an enrichment of down-regulated clusters.

## RESULTS

### Non-random Clustering of topologically Co-Regulated Genes

We first sought to define domains with concordant response to DNA topological stress in the form of gene clusters of contiguous GRO values (Figures 1A,B and Methods). In total there were 116 clusters with more than 7 genes and 180 clusters containing 6 or more genes, which were deemed highly significant on the basis of a permutation test (Figure 1C, Methods). Of these significantly long (>=7 genes) clusters, 50 comprised exclusively up-regulated genes and 66 exclusively down-regulated ones (median number of genes=8 for both types, Supplementary File 2). In total, the clusters comprised 1074 genes (~20% of the total).

Given that we measure topological stress in transient heat shock conditions, we wanted to see if the clustering effect we observe could be attributed to the temperature shift. We employed an identical clustering approach in gene expression profiles obtained upon heat shock stress conditions as published in a landmark paper (31) for both transient (20min) and prolonged (80min) heat-shock (HS). Even though there is some degree of overlap the numbers and sizes of the transient and prolonged HS-stress clusters are very small (19 and 12 clusters respectively, comprising less than 6% of the total genes, Supplementary Figure 2). Hence, the observed strong clustering tendency appears to be a characteristic property of the topologically induced/suppressed genes.

Genes belonging to the topological stress-induced genes showed a significant tendency to be co-regulated. By analyzing weighted gene expression correlations based on the largest compendium of gene expression experiments in yeast (32), we found topologically induced clusters to be have significantly greater adjusted correlations scores (ACS) compared to a random selection of gene clusters (Figure 1D). Based on the way they were defined, we chose to refer to them as “Topologically co-regulated gene clusters” (TCGC) and went on to characterize them in terms of various properties.

### Positional Preferences of Topologically Co-regulated Gene Clusters in linear chromosomes

The distribution of TCGC (Figure 2A) suggests a non-random localization along the genome. Up-regulated gene clusters tend to be found towards the outer boundaries of linear chromosomes, while down-regulated ones show a tendency for their center, often in close proximity to the centromeres. In some cases, clusters appear to assemble in super-clusters as in the case of the right arm of chromosome 12 or the left arms of chromosomes 6 and 7. A straight-forward analysis of TCGC distance from the centromeres showed statistically significant opposing preferences for the up– and down-regulated gene clusters to be located away from and close to centromeres respectively (p<=0.05, Supplementary Figure 4).

**Figure 2.**
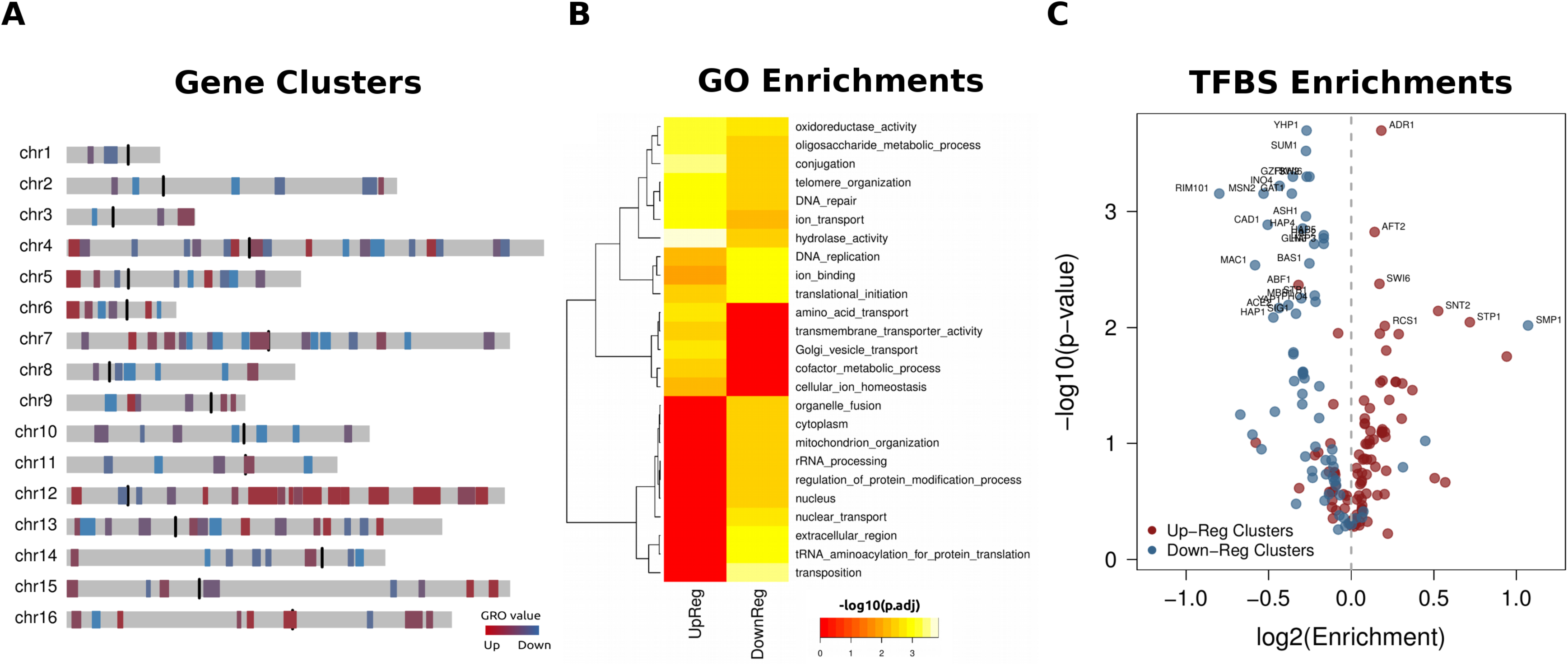
Functional Enrichment and Regulatory Modes in Topologically Coregulated Clusters. **A.** Distribution of 116 topologically co-regulated gene clusters (TCGC) in the yeast genome. **B.** GO term enrichment heatmap of TCGC of both types. Enrichments were calculated based on a modified Gene Set Enrichment Analysis (26). Only GO terms with an adjusted p-value<=0.05 (at 5% FDR) for at least one of the two TCGC types are reported. **C.** Volcano plot showing enrichments of transcription factor binding sites for 102 different transcriptional regulators compiled by (27). Enrichments are shown as log2 based observed/expected ratios. Values >0 indicate enrichment and values <0 indicate depletion (see Methods). P-values correspond to 1000 bootstraps for each transcriptional regulator.

The process of DNA replication is tightly connected to DNA supercoiling. We sought to examine differences in the positions of TCGC compared to DNA replication origins (ARS). We found down-regulated clusters to be preferentially located away from DNA replication origins (ARS) (Mann-Whitney U-test p<=0.0003 compared to a random set of clusters). Even though this may be related to a lack of ARS sites in close proximity to the centromeres, the great discrepancy between down– and up-regulated TCGC to overlap DNA replication origins (Fisher’s test, p=0.000381, OR=0.175) is characteristic of strong opposite positional preferences between the two types of clusters. This avoidance may be explained on the basis of an optimization strategy throughout the course of evolution, as genes that are more severely affected by topological stress will tend to be located far from DNA replication origins.

### Opposing Functional and Regulatory Preferences in different types of TCGC

TopoII is essential for yeast cells and its prolonged deactivation is bound to cause a general shutdown of cellular activity. The fact, however, that a significant proportion of yeast genes respond to its transient deactivation with increased transcription levels indicates the existence of a positive effect for a subset of cellular functions. In a previous study (18) we have shown that the regulatory and functional properties of the topo-affected genes is radically different from those induced or repressed upon heat shock. In fact, the stress imposed by the deactivation of topoII is very particular, when compared to a number of nutrient, chemical or physical stresses (Supplementary Figure 3). This prompted for a detailed functional analysis of the topologically deduced gene clusters.

A functional enrichment analysis at the level of Gene Ontology (Figure 2B) shows extensive differences between the two types of TCGC, a fact indicative of their nuclear compartmentalization being echoed in their functional roles. Three main clusters are apparent: a) Functions enriched in both types of clusters include secondary metabolism and DNA maintenance. b) GO terms that are enriched in up-regulated clusters and depleted in down-regulated ones, represent functions related to cellular transport, the metabolism of co-factors and general stress response. c) Down-regulated-specific GO terms contain basic cellular functions associated with RNA transcription and processing, translation and the nuclear environment. A general pattern suggests that the localization of clusters among chromosomes is also reflected in their functions with up-regulated gene clusters being mostly enriched in peripheral functions, unrelated to the core nuclear processes, the opposite being the main characteristic of genes within down-regulated clusters.

Figure 2C highlights the transcription factors whose binding sites are found more or less frequently than expected by chance for both types of TCGC. Down-regulated gene clusters tend to be mostly depleted of TFBS, partly explained by the fact that they are enriched in constitutively expressed genes and thus subject to less complex regulation. From a previous analysis at the level of genes on the same dataset we know down-regulated genes to be enriched in essential functions, with constant expression levels and mostly depleted of TATA-boxes (18). Up-regulated TCGC, on the other hand, show the exact opposite pattern, with the great majority of the TFBS being enriched, a fact indicative of more complex regulation, with significant enrichments for factors related to chromatin structure, DNA surveillance and amino acid transport (see Supplementary Table 1 and discussion). This positional-functional compartmentalization is also reflected on a number of structural attributes of these clusters, discussed in the following.

### Gene Spacing and Directionality of Transcription in TCGC

During transcription, DNA torsional stress accumulates with different sign ahead of and behind the gene’s transcription start site. This makes the size of both the gene and the preceding intergenic spacer, as well as the relative direction of transcription in relation to adjacent genes highly relevant for the dissipation of topological tension. The effect of topoII deactivation has been shown to be generally independent from the size of the majority of yeast genes (33), but it is strongly inhibitory in the case of long transcripts (34). The situation is very different when one looks, instead, into the surrounding intergenic space. When we ranked the complete set of yeast genes according to their GRO values and plotted them against the mean size of intergenic spacers we found a a clear positive correlation (p-value<=10^-12^) between the log-size of the intergenic regions and the GRO value. This is highly indicative of transcription-induced topological stress being more readily dissipated in genes with long upstream (and downstream) regions (Supplementary Figure 6). It is worth noticing, that this is another characteristic property of topologically-induced gene clusters. When comparing the intergenic spacer sizes of up– vs down-regulated gene clusters a significant difference was found for TCGC (t.test p-value <=10^-6^, Heat-shock clusters p-value=0.09).

The association between DNA topology and structural genomic features is expected to be more pronounced in the series of adjacent genes with similar GRO values. In order to study the effect of intergenic space in co-regulated gene clusters, we employed a more relaxed criterion in the definition. We thus obtained all possible arrays of 7 contiguous genes, ranked them according to their mean GRO value and kept the top and bottom 200 non-overlapping such arrays as up-regulated and down-regulated clusters. These contained the complete set of our TCGC but also a number of additional gene clusters that showed consistent behaviour in their response to topological stress, although not entirely positive or negative in terms of GRO value. We then expanded these clusters on either side in order to comprise 11 genes each (see Methods for details) and compared the average intergenic space along them as shown in Figure 3A. Up-regulated clusters showed intergenic regions of significantly increased size compared to the genomic average (which is about 660bp), an increase that, moreover, appeared to be inflated towards the central genes in the cluster. Genes in down-regulated clusters were, on the other hand, flanked by much shorter intergenic regions and did so consistently, with little fluctuation. This striking discrepancy is not observed in the case of heat-shock induced clusters (see Supplementary Figure 7A). Besides their reduced potential for resolving topological stress, shorter intergenic regions provide shorter available genomic space for transcription factors, which may account for the marked under-representation of TFBS in down-regulated gene clusters (Supplementary Figure 8B).

**Figure 3.**
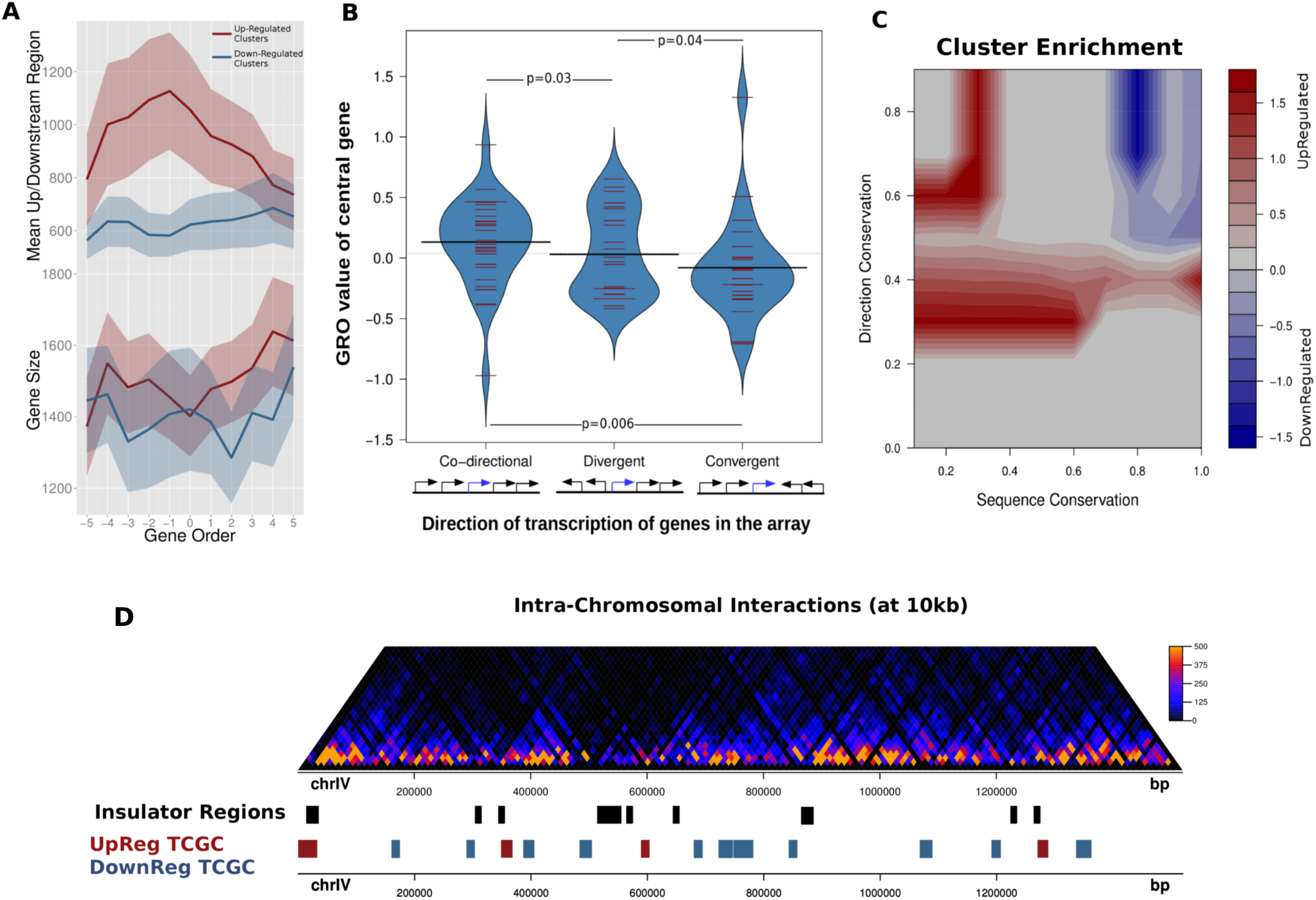
Topologically co-regulated genes share distinct preferences for intergenic space and transcription directionality. **A.** Top, mean intergenic region length for clusters of 11 consecutive genes. Each line corresponds to the mean values calculated for the top/bottom 200 clusters based on the central 7 GRO values (see Methods for details). Bottom, same analysis for gene size. Shaded bands correspond to 95% confidence intervals. **B.** Distribution of GRO values of genes lying in the center of 5-cluster genes with different directionality patterns defined on the basis of transcriptional direction (N co-directional=36, N divergent=29, N convergent=25). P-values calculated on the basis of a Mann-Whitney U test. **C.** Contour heatmap of enrichment of different types of TCGC in areas defined by mean sequence conservation (as above, x-axis) and a transcriptional direction index (y-axis) defined as the proportion of genes retaining relative gene position and directionality in two closely related species. Enrichments were calculated as log2(ratios) of proportion of up-regulated/down-regulated clusters having values in a 10x10 value grid (see Methods). **D.** Intra-chromosomal interactions map of chromosome IV and corresponding domains of insulation (in black). Up-regulated clusters (red) show a significantly increased tendency to be located in proximity to insulation regions, when compared to down-regulated ones.

Figure 3A is strongly indicative of the impact of genomic architecture on the maintenance of topological equilibrium in the nucleus. Genes flanked by shorter intergenic spacers will be more prone to the accumulation of supercoiling on either side of the transcription bubble and are therefore expected to be more sensitive to the lack of topoII, while genes that allow for the dissipation of topological strain into longer, untranscribed, nearby regions are predictably more resilient.

Synergistic effects between neighbouring genes may be accentuated by the directionality of transcription of consecutive genes. Gene clusters with more “streamlined” directionality patterns are expected to be able to accommodate DNA supercoiling in a more effective manner, using alternating positive and negative supercoiling to “propel” transcription. In order to test this hypothesis, we searched our gene cluster dataset for specific patterns of gene directionality. We split clusters in three categories depending on whether the central gene in the cluster a) formed part of a series of co-directional transcriptional units or b) was belonging to a pair of divergently or c) convergently transcribed genes. We then compared the GRO values of the central gene in each category. The results (Figure 3B, Supplementary Figure 9) are indicative of a mild, yet significant association between gene directionality patterns and response to topoII deactivation. Genes lying midway in clusters of co-directional transcription have in general higher GRO values, while genes belonging to convergent pairs have difficulty in dealing with topological tension. Divergently transcribed genes lie somewhere in the middle in terms of sensitivity as reflected in their average GRO values. Again, this appears to be a distinctive property of TCGS (compare with heat-shock clusters, Supplementary Figure 7B).

### Different Conservation Constraints in TCGC

In order to investigate how the properties described above may be constrained through evolution, we performed an analysis of conservation at two levels. First, we analyzed the mean sequence conservation per cluster as aggregate phastCons scores (29), obtained from a genome-wide alignment of six *Saccharomyces* species (35). Average sequence conservation (excluding intergenic space) was negatively correlated with the mean GRO value for the 116 TCGC (p<=0.01), confirming that down-regulated clusters are significantly more constrained in terms of sequence conservation (Supplementary Figure 8A). Increased conservation for down-regulated gene clusters doesn’t come as a surprise given their functional preferences described in previous sections. Genes in up-regulated clusters on the other hand appear to be under more moderate sequence constraint, a fact which could be indicative of their less essential role, or their more recent acquisition through gene duplication (36).

We next turned to more complex conservational features that also take into account synteny relationships, reflected upon the position and transcriptional direction of genes in related species. We made use of data from the Yeast Gene Order Browser (YGOB; http://wolfe.gen.tcd.ie/ygob) (30) that contains a detailed catalog of orthologous genes between a number of yeast species. We collected all orthologous gene pairs between *S.cerevisiae* and two of its closest species in the *sensu strict*o complex, *S. paradoxus* and *S. mikatae*. We analyzed them separately for up– and down-regulated clusters by calculating a simple measure of “directional conservation” (Methods). Given that syntenic regions are by definition under sequence constraint we were not surprised to see that genes in down-regulated clusters were characterized by both high sequence and directional conservation as may be seen in Figure 3C. What was rather interesting was the corresponding position of genes in up-regulated clusters in the same two-dimensional constraint space. While we already knew that sequence constraints were more relaxed in these regions, we found a significant proportion of genes with high values of directional conservation, suggesting that up-regulated gene clusters tend to maintain the directionality patterns even under milder sequence constraints. It thus seems, that keeping a co-directional gene layout confers a relative advantage to genomic regions that are otherwise less conserved in terms of sequence.

### Topologically Co-regulated Gene Clusters associate with different components of the three-dimensional genome structure

The eukaryotic nucleus is organized in three-dimensions, where chromosomes interact in space forming intra and inter-chromosomal domains (37) and which largely affect the processes of genome replication and transcription. Even though, the three-dimensional organization of yeasts does not share the complexity of higher eukaryotes with topologically-associated domains (TAD) and nuclear compartments, it maintains aspects of organization such as “globules” that represent regions of increased intra-chromosomal interactions (23,38). Having observed strong positional preferences of TCGC in linear dimension, we went on to examine whether these may be reflected in the higher-order three-dimensional structure of the genome. We used intrachromosomal interaction data from a 3C experiment (23) to define TAD-like globules separated by insulating regions (see Methods). We found down-regulated gene clusters to be predominantly occupying regions spanning the globules, while up-regulated ones were found to be significantly more proximal to insulating boundaries (Figure 3D, Fisher’s test p-value=0.0182, Supplementary Figure 10). This propensity is indicative of a preference of clusters that are positively affected by topological stress to occupy less entangled and more “open” parts of the chromatin and fits well with the rest of their characteristic properties described up to now.

Differences in the distribution of TCGC in the three-dimensional nucleus were also picked up after an analysis at a higher level using a partition of the yeast genome in chromosomal networks (25) (see Methods). We found down-regulated TCGC to be preferentially located in the center of the nucleus, described in the model of (25) as an extensive “community” of pericentromeric interchromosomal interactions. Up-regulated ones, on the other hand, were mostly found enriched in the periphery, which is constituted by the subtelomeric regions and the right arm of chromosome 12 (Supplementary Figure 5).

## DISCUSSION

The existence of clusters of topologically co-regulated genes (TCGC) implies that eukaryotic genes may be synergistically orchestrated in gene neighborhoods with particular characteristics. By persistently analyzing the defined gene clusters at various levels, we were able to outline an overarching pattern, according to which the yeast genome may be broadly divided in two compartments that have, in time, assumed radically different architectures and operational roles. Down-regulated clusters, preferentially located towards the center of the nucleus, consisted of highly conserved genes associated with essential functions. These are expectedly shut-down by the accumulation of supercoiling in the process of a general suspension of topologically “expensive” processes such as the transcription of rRNA. Up-regulated clusters, on the contrary, comprise stress-responsive genes, whose functions may be required to dampen or even reverse topological stress. Their structural organization, with long intergenic spacers and gene transcriptional co-directionality may further enable these areas of the genome to even harness DNA supercoiling in order to achieve increased transcription levels.

Such compelling disparity at all studied levels points towards a general pattern of genome architecture. This very much resembles an urbanization process, that has over evolution demarcated an “old-town” at the centromeric part of the nucleus, formed by tightly crammed ancient genes and a “suburban genome” at the chromosomal outskirts, where newly acquired genes occupy greater spaces with an ordered directionality that resembles tract housing (Figure 4). This “Genome Urbanization” is echoed in various genomic features that we have discussed in the context of TCGC. When looking at the sequence conservation of genes as a function of their distance from the centromere we find a weak negative correlation, with the 5% most distant genes being significantly less conserved than the 5% most proximal (n=638, t.test p-value=0.005). Similar discrepancy is observed when looking at the intergenic space length (n=508, t.test p-value<10^-6^). It thus appears that the division of the genome in domains with specific “architectural” characteristics may well extend beyond DNA topology. Our findings indicate that the Genome Urbanization scheme is likely a general feature, that allows the nucleus to dissipate DNA topological stress more effectively, but whose functions are likely to extend to gene functionality (39), regulation programs (16,40) and genome evolution (41).

**Figure 4.**
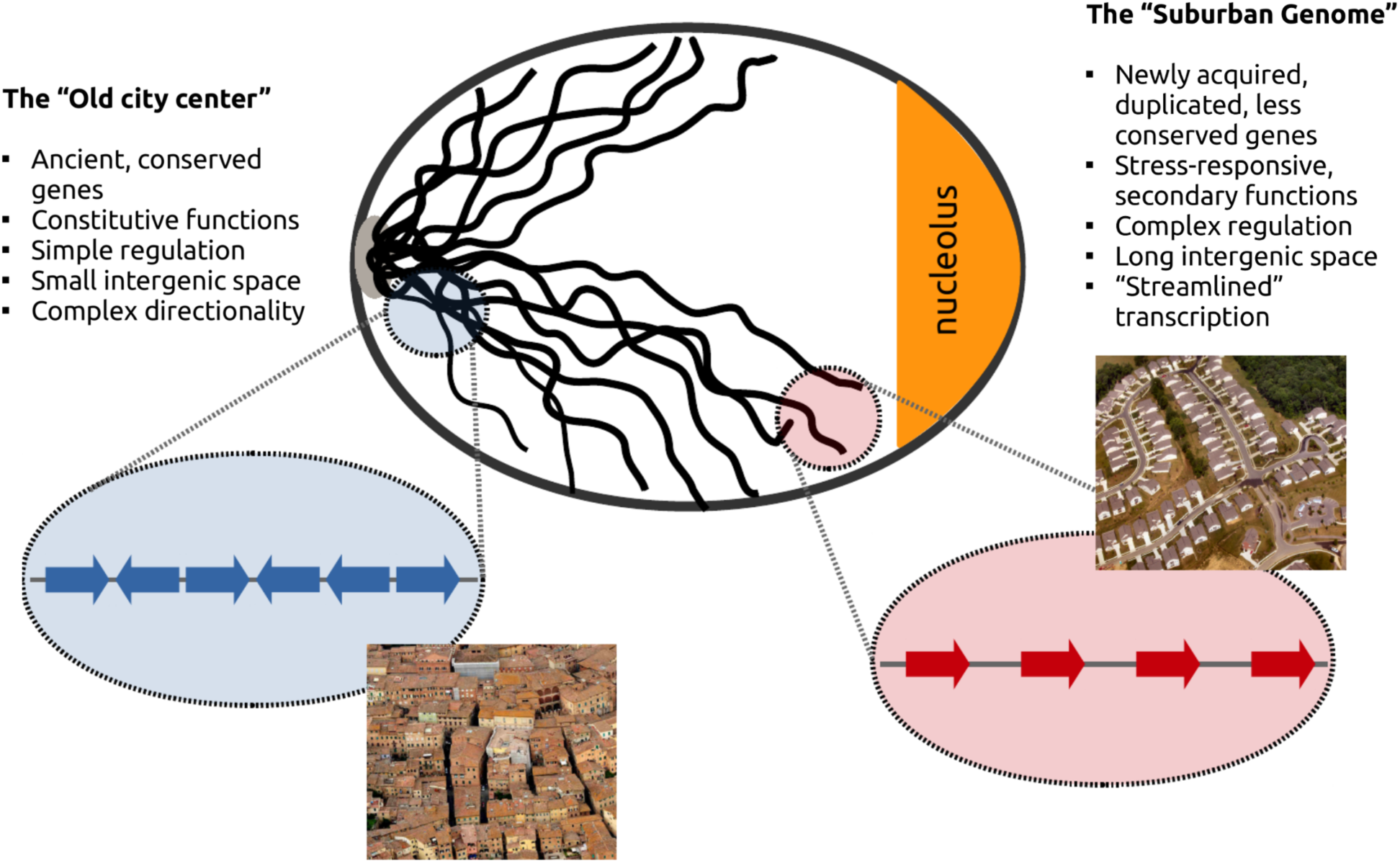
Genome Urbanization. Positional preferences of topologically co-regulated genes reflect structural, regulatory and functional compartmentalization. Genome Urbanization in *S. cerevisiae*. A schematic of the yeast interphase nucleus is shown based on the Rabl configuration (45). Pericentromeric regions correspond to what we call the “Old city center” with enrichment in gene clusters down-regulated under topoII deactivation. The genome in these areas may be compared to the crammed houses of a medieval town separated by narrow, intertwined alleys. Genes in the “old town” are more conserved, associated with essential functions and located within tighter genomic spaces with fewer transcription factor binding sites and entangled directionality. Genomic regions at the nuclear periphery are resembling a “suburban landscape” where more recently acquired (and less conserved) genes are spaced in co-directional operon-like arrays, separated by longer intergenic sequences, reminiscent of the tract housing of modern city suburbia.

A particularly important element to consider is that of transcriptional plasticity. The over-representation of stress responsive genes in up-regulated clusters points towards an organization of the genome, in which genes that need to readily modulate their expression levels according to environmental conditions are preferentially located in particular genomic “niches”. Recent works have provided interesting links between plasticity and genomic features that resemble the ones we find to be hallmarks of the “suburban genome”, namely non-essentiality, complex regulation and gene duplication (42). The size of the intergenic space between genes has also been shown to widely shape expression variability (43).

The concept of “Genome Urbanization” may extend to more complex eukaryotes, albeit not in a straight-forward manner. The size, gene density and evolutionary dynamics of the unicellular *S. cerevisiae* make the delineation of domains more clear-cut, while the complexity of gene-sparse genomes from multicellular organisms with the requirements for spatio-temporal expression patterns is bound to be reflected upon a more entangled genome architecture (44). The advent of new experimental approaches for the study of genome conformation in three dimensions provides a solid framework for testable hypotheses that will deepen our understanding of the evolution of genome organization.

## ACKNOWLEDGEMENTS

The authors thank Despoina Alexandraki and Babis Spilianakis for a critical reading of an initial draft of the manuscript and Roderic Guigo for fruitful discussion on the presented aspects.

